# GPS2 regulates mitochondria biogenesis via mitochondrial retrograde signaling and chromatin remodeling of nuclear-encoded mitochondrial genes

**DOI:** 10.1101/162297

**Authors:** Maria Dafne Cardamone, Bogdan Tanasa, Carly Cederquist, Jiawen Huang, Kiana Mahdaviani, Wembo Li, Michael G. Rosenfeld, Marc Liesa, Valentina Perissi

## Abstract

As most of the mitochondrial proteome is encoded in the nucleus, mitochondrial functions critically depend on nuclear gene expression and bidirectional mito-nuclear communication. However, mitochondria-to-nucleus communication pathways are incompletely understood. Here, we identify G-Protein Pathway Suppressor 2 (GPS2) as a mediator of mitochondrial retrograde signaling and a key transcriptional activator of nuclear-encoded mitochondrial genes in mammals. GPS2 regulated translocation from mitochondria to nucleus is essential for the transcriptional activation of the nuclear stress response to mitochondrial depolarization and for supporting basal mitochondrial biogenesis in differentiating adipocytes and in brown adipose tissue from mice. In the nucleus, GPS2 recruitment to target gene promoters regulates histone H3K9 demethylation and RNA Polymerase II (POL2) activation through inhibition of Ubc13-mediated ubiquitination. Together, these findings reveal an unexpected layer of regulation of mitochondrial gene transcription as they uncover a novel mitochondria-nuclear communication pathway.

## Introduction

Normal mitochondrial function is vital to the well being of eukaryotic cells. Beyond ATP production, mitochondria generate key metabolites and biochemical signals central to apoptotic and metabolic regulatory pathways. Conversely, defects in mitochondrial functions have been associated with many human diseases, including not only primary inherited mitochondrial disorders, but also insulin resistance and Type 2 diabetes (T2D) (Altshuler-Keylin and Kajimura, 2017; Gorman et al., 2016; Hu and Liu, 2011; Kusminski and Scherer, 2012; Newsholme et al., 2012; Patti and Corvera, 2010; Wallace, 2013). Even though mitochondria are semi-independent organelles containing multiple copies of their own genome, most of the mitochondrial proteome is encoded by nuclear genes and, thus, regulated by nuclear transcription factors (TFs) and associated cofactors including PGC1α, NRF1, GABPα, ATF5, ERRα and others (Brenmoehl and Hoeflich, 2013; Fiorese et al., 2016; Hock and Kralli, 2009; Scarpulla et al., 2012; Whelan and Zuckerbraun, 2013; Wu et al., 1999). Thus, the biogenesis of mitochondria and the maintenance of mitochondrial homeostasis critically depend on nuclear transcription and2 on the presence of signaling pathways that inform the nucleus of mitochondrial dysfunctions and changes in cellular metabolism (Finley and Haigis, 2009; Guha and Avadhani, 2013; Hu and Liu, 2011; Kotiadis et al., 2014; Vafai and Mootha, 2012; Yun and Finkel, 2014). This is partly achieved through regulation of nuclear TFs and cofactors via cytosolic mediators (i.e. Ca2+, ROS, NAD/NADH ratio)(Bohovych and Khalimonchuk, 2016; Hock and Kralli, 2009; Quiros et al., 2016; Scarpulla, 2012; Scarpulla et al., 2012; Wu et al., 1999). However, much is still unknown about mitochondria-to-nuclear signaling pathways and about the physiological effects of impaired communication networks in the development of diseases (Chandel, 2015; Finley and Haigis, 2009; Guha and Avadhani, 2013; Vafai and Mootha, 2012).

A direct mitochondrial retrograde response pathway was first described in response to mtDNA depletion in *S. Cerevisiae*. Key players in this pathway are the DNA transcription factors Rtg1-Rtg3, and the regulatory factor Rtg2, which drives Rtg1-3 translocation to the nucleus in response to changes in ATP availability and mitochondrial membrane potential (MMP)(Jazwinski and Kriete, 2012; Jia et al., 1997; Liao and Butow, 1993; Rothermel et al., 1995; Rothermel et al., 1997; Sekito et al., 2000). Partially overlapping functions of Rtg1/Rtg3 with mammalian transcription factors (such as FOXOs, NFkB, ERα and Myc) speak to the functional conservation of this pathway, even in absence of clear mammalian orthologues of the yeast *Rtg* genes (Germain, 2016; Jazwinski, 2013; Quiros et al., 2016). Investigating mito-nuclear communication strategies in response to proteotoxic stresses (UPR^mt^) in worms also led to the characterization of a transcriptional response program coordinated by transcription factor ATFS-1 upon nuclear translocation, further supporting the notion that direct communication strategies might be employed across species (Lin et al., 2016; Nargund et al., 2015; Nargund et al., 2012). However, it is currently unknown whether direct mediators of retrograde signaling, yet to be identified, exist in mammals.

GPS2 is a multi-functional protein that recently emerged as an important regulator of inflammation and lipid metabolism, processes tightly linked to mitochondrial functions (Cardamone et al., 2012; Cardamone et al., 2014; Jakobsson et al., 2009; Toubal et al., 2013; Venteclef et al., 2010). The nuclear GPS2 protein content is tightly controlled3 through protein stabilization/degradation (Huang et al., 2015) and GPS2 deficiency in mice is embryonic lethal (Guo et al., 2014), indicating the physiological importance of GPS2. Previous work from us and other laboratories indicate that nuclear GPS2 acts as a transcriptional cofactor playing a dual role as corepressor and coactivator for a number of transcription factors (Cardamone et al., 2014; Cheng and Kao, 2009; Guo et al., 2014; Jakobsson et al., 2009; Peng et al., 2001; Venteclef et al., 2010; Zhang et al., 2008). Furthermore, in addition to the nuclear functions, we recently reported a number of distinct, but possibly coordinated, functions for GPS2 in the cytosol as regulator of insulin signaling and pro-inflammatory response pathways (Cardamone et al., 2012; Cederquist et al., 2017; Lentucci et al., 2016). While the molecular mechanisms underlying GPS2 genomic and non-genomic functions have not been fully elucidated, our work suggests that both nuclear and cytosolic functions are linked to GPS2-mediated inhibition of lysine 63 (K63) ubiquitination events (Cardamone et al., 2012; Cardamone et al., 2014; Cederquist et al., 2017; Lentucci et al., 2016). This raises a number of important questions regarding GPS2 function and regulation in the cell. First, does GPS2 shuttles between different intracellular locations? Second, is its extra-nuclear residence important for informing and modulating GPS2-mediated transcriptional regulation? Third, do the opposing functions of GPS2 and the K63 ubiquitin conjugating enzyme Ubc13 serve a key role for the coordinated regulation of metabolic and inflammatory pathways? The current study addresses these questions by demonstrating that GPS2 regulated translocation from mitochondria to nucleus represents a novel mitochondria retrograde signaling pathway, which is required for promoting the transcriptional reprogramming induced by mitochondria depolarization and for supporting mitochondria biogenesis during adipocyte differentiation. Mechanistically, our results indicate that GPS2-mediated inhibition of Ubc13 activity on target promoters is required for promoting the dimethylation of repressive histone mark H3K9me3 and the activation of POL2-mediated transcription, thus suggesting that modulation of non-proteolytic ubiquitination represents an unappreciated hub for integrating metabolic pathways modulating the balance between lipid storage and energy expenditure in the adipose tissue.

## Results

### Dual mitochondrial/nuclear localization of transcriptional cofactor GPS2

To better elucidate GPS2 complex role within the cell, we first sought out a more detailed description of its subcellular localization. In agreement with its proposed functions, GPS2 was previously reported in both the nuclear and cytosolic compartments (Cardamone et al., 2012). A closer inspection of GPS2 intracellular localization by immunofluorescence revealed that the extra-nuclear staining was both diffused in the cytosol as well as concentrated in specific structures. In agreement with *in silico* prediction analyses using TargetP1.1 and Mitoprot (Claros and Vincens, 1996; Nielsen et al., 1997), co-staining with mitochondrial markers mtHSP70 and ATP5B indicated that the punctate cytosolic staining corresponded to GPS2 localization to mitochondria in both murine 3T3-L1 and human Hela cells (**Fig. 1A**). Specificity of the mitochondrial staining was confirmed in 3T3-L1 cells by GPS2 downregulation with two independent siRNA (**Supplemental Fig. S1A** and **S1B**). Conversely, the expression of a HA-GPS2 fusion construct resulted in extensive HA mitochondrial staining in both 3T3-L1 and Hela cells (**Supplemental Fig. S1C**), confirming mitochondrial targeting of the fusion protein. Mitochondria localization of endogenous GPS2 was also confirmed by immunogold labeling coupled with electron microscopy in Hela cells (**Supplemental Fig. S1D)**. These results together identify GPS2 as a previously unrecognized mitochondrial protein.

**Figure 1.**
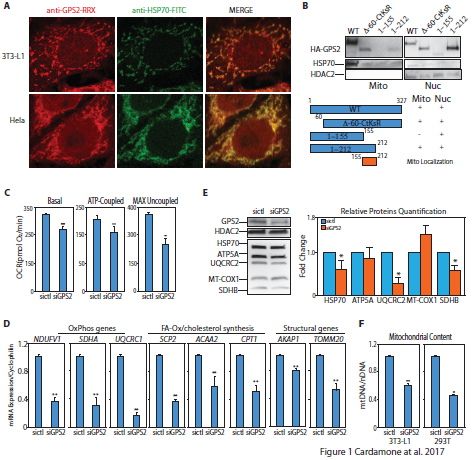
GPS2 localizes to mitochondria and regulates mitochondrial function through nuclear-encoded mitochondrial gene expression. **(A)**Mitochondria localization of endogenous GPS2 by IHF staining in murine 3T3-L1 and human Hela cells. Co-staining of endogenous mtHSP70 serves as a mitochondrial marker. **(B)**Identification of minimal mitochondria localization domain by western blotting of fractionated extracts in 293T cells transfected with full length GPS2 (wt), GPS2 with the first 60 aa deleted (?60-CtKsR), GPS2 fragment 1-155 and 1-212. WBs for HDAC2 and mtHSP70 respectively serve as control for nuclear (Nuc) and mitochondrial (Mito) extracts. **(D)**Decreased respiration and oxygen consumption, as measured by Seahorse, in 3T3-L1 cells upon GPS2 transient downregulation. **(E)**RT-qPCR analysis of the expression of different classes of nuclear-encoded mitochondrial genes in 3T3-L1 cells transfected with siCTL or siGPS2. **(F)**Western blot analysis of mitochondrial proteins expression with relative quantification (siGPS2 vs siCTL). **(G)**Mitochondrial content measured by PCR as a ratio of mtDNA/nDNA in WT and GPS2-KD 3T3-L1 and 293T cells. Results in **(C), (D),** and **(F)** represent the sample mean of three independent experiments; error bars represent s.e.m. (n=3). *indicate p-value<0.05; **\*\*** indicate p-value<0.01.

Next, we wished to define the domain responsible for GPS2 mitochondrial localization. Because inspection of the GPS2 sequence did not reveal any classic mitochondria localization signal (MLS), we overexpressed different GPS2 deletion constructs and verified the protein subcellular localization by fractionation and WB. Because we had previously shown that N-terminal deletions lead to protein degradation, deletion of the first 60 aa was tested in association with point-specific mutations of the lysines targeted by protein ubiquitination (K254, 300, 327A)(Huang et al., 2015). In this stabilized form, deletion of the N-terminus did not affect mitochondrial localization, confirming the absence of an N-terminal mitochondrial import sequence. Likewise, the C’terminus domain was not required for mitochondrial localization (**Fig. 1B**). Deletion of aa.155-327, instead, severely impaired the expression of HA-GPS2 in the mitochondria, indicating that mitochondrial localization of GPS2 is likely driven by an internal import signal between aa. 155 and 212 (**Fig. 1B**).

### GPS2 regulate mitochondrial function through transcriptional regulation of nuclear-encoded (neMITO) genes

To investigate the physiological relevance of GPS2 in the regulation of mitochondrial functions, we measured mitochondrial respiratory capacity in intact cells depleted of GPS2 by siRNA-mediated downregulation (GPS2-KD). For these experiments, we transiently downregulated GPS2 in either 3T3-L1 cells, using one of the murine siRNA validated above, or Hela cells, using a distinct human siRNA that was previously used to characterize GPS2 in 293T cells (Cardamone et al., 2012). In both systems GPS2 downregulation caused a significant decreases in maximal respiratory capacity and ATP-linked respiration (**Fig. 1C and Supplemental Fig. S1E**). Because of GPS2 well-described role in transcriptional regulation, we asked whether the observed lower cellular respiration in absence of GPS2 was due to a defect in the expression of mitochondrial genes. In agreement with this hypothesis, GPS2 was consistently required for the expression of a variety of nuclear-encoded mitochondrial (neMITO) genes selected across different functional classes (**Fig. 1D,** for validation with distinct siRNA see **Supplemental Fig. S1F**). In contrast, the expression of mtDNA-encoded genes was not affected by GPS2 downregulation (**Supplemental Fig. S1G**). These results were confirmed by corresponding changes in protein levels of representative nuclear-encoded and mitochondrial-encoded enzymes (**Fig. 1E**). Also, rescue experiments with HA-GPS2 restored the expression of neMITO genes, confirming the on-target specificity of this effect (**Supplemental Fig. S1H**). In addition, GPS2 downregulation caused a significant reduction in mitochondrial DNA copy number in both 3T3-L1 and 293T cells (**Fig. 1F**). This suggested that a decline in the amount of mitochondria, possibly caused by a global reduction in mitochondrial gene expression, rather than a specific defect in electron transport chain activity, might be responsible for the observed reduced respiration capacity of GPS2-KD cells.

To further address the relevance of GPS2 as a global regulator of mitochondrial gene expression, we performed genome-wide RNAseq profiling of 3T3-L1 WT and GPS2-KD cells. Considering the full transcriptome, we identified 1086 Differentially Expressed Genes (DEGs) upon GPS2 downregulation, including 439 genes down-regulated and 647 genes up-regulated (**Fig. 2A** and **Supplemental Table 1**). Mitochondrial GO terms were among the most significantly enriched categories among down-regulated genes, whereas up-regulated genes associated with non-mitochondrial terms (**Fig. 2A,** EnrichR analysis of GO Cellular Pathways (Chen et al., 2013)). In agreement with the significant enrichment in mitochondrial terms observed among downregulated genes, approximately 20% of the downregulated genes compared to 10% of the upregulated were mitochondrial genes (**Fig. 2A**). Furthermore, only downregulated genes were enriched for canonical mitochondrial pathways like “TCA cycle” and “Electron transport chain” (**Fig. 2A,** EnrichR analysis of WikiPathways (Chen et al., 2013)). Upregulated mitochondrial genes were more loosely associated with “Interleukin signaling pathways”, “SREBP signaling” and “NFkB signaling” (**Fig. 2A)**, suggesting they could represent a secondary response to mitochondrial dysfunction or to the upregulation of AKT and pro-inflammatory pathways in the cytosol. Accordingly, total upregulated genes were enriched for “Focal adhesion”, “Regulation of actin cytoskeleton” and “PI3K/AKT pathway” (**Supplemental Fig. S2A**), as expected based on GPS2 regulating insulin signaling through AKT activation in the adipose tissue (Cederquist et al., 2017). Downregulated genes were instead enriched for “Cell cycle”, “Meiosis” and “Glutathione metabolism” functional pathways (**Supplemental Fig. S2A,** EnrichR analysis of KEGG Pathways (Chen et al., 2013)).

**Figure 2.**
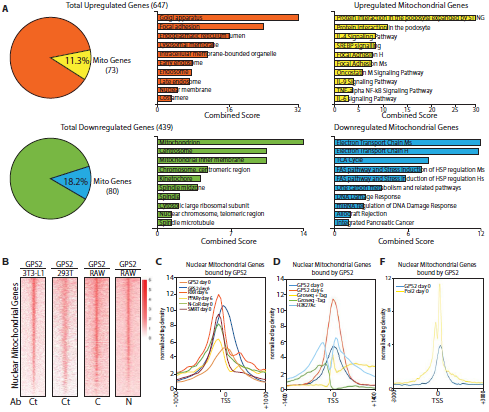
GPS2 binds to the core proximal promoter of nuclear-encoded mitochondrial genes to regulate their transcriptional activation. **(A)**DEGs genes between WT and GPS2-KD 3T3-L1 cells as identified by RNAseq (two independent biological replicates, FDR<0.05). Mitochondrial genes are enriched among downregulated genes. Central plot represents the 10 most significant GO terms associated with total upregulated (orange) and downregulated (green) genes (EnrichR, GO cellular component). Right plot represents the 10 most significant pathways associated with upregulated (yellow) and downregulated (blue) mitochondrial genes (EnrichR, WikiPathways). GO terms and Pathways are listed based on a combined score between p-value and Z-score (EnrichR). **(B)**Heatmaps of GPS2 binding on mitochondrial genes by ChIPseq. GPS2 binding profile in 3T3-L1 and 293T cells were generated using an antibody against aa 307-327 (Ct) (GSE57779 and GSE35197). GPS2 binding in RAW macrophages was generated by Fan and colleagues (Nature Medicine, 2016) using distinct antibodies against the N terminus (N) and C terminus (C) of GPS2 (GSE66774). Represented in the heatmaps are all mitochondrial genes (2086 murine genes and 2296 human genes). **(C)**Tag density plot of GPS2 binding on mitochondrial genes compared to the binding for RXR, PPARγ, NCoR and SMRT (GSE13511 and Array Express E-MTAB-1031) in differentiating 3T3-L1 (profiled at either day 0 or day 6 of differentiation). **(D)**Tag density plots of GPS2 peaks on mitochondrial genes in 3T3-L1 cells compared to GROseq tags (GSE56745) and ChIPseq tags for H3K27Ac (GSE36965) reveal GPS2 binding over the transcriptional start site (TSS) area. **(E)**Tag density plot of GPS2 binding on mitochondrial genes compared to POL2 binding (GSE13511) in at al. 2017

Notably, comparable results were observed in a previously published transcriptomic analysis in human 293T cells (Cardamone et al., 2012)(**Supplemental Table 2**). The transcriptional signature of GPS2-deleted B cells also included a similar enrichment in the downregulation of genes associated with mitochondrial functions (Lentucci et al., 2016). Thus, profiling of GPS2 transcriptional signature in different cell models confirms that GPS2 is broadly required for the activation of mitochondrial gene expression and reveal that its downregulation is associated with significant reprogramming of cell growth and metabolism.

As these results suggested that GPS2 might be regulating mitochondrial gene expression as a nuclear cofactor, we next re-analyzed previously published GPS2 ChIPseq datasets (Cardamone et al., 2012; Cardamone et al., 2014; Fan et al., 2016) by focusing on mitochondrial genes. Strikingly, GPS2 binding was observed on most mitochondrial genes in each cell model, including macrophages, adipocytes and embryonic kidney cells using three distinct antibodies **(Fig. 2B)**. In each of these models, GPS2 binding sites genome-wide had been previously mapped to both promoter and distal regulatory regions on distinct and overlapping target gene programs (Cardamone et al., 2012; Cardamone et al., 2014; Fan et al., 2016). In contrast, GPS2 binding to neMITO genes was found concentrated on promoter areas, with the enrichment in GPS2 binding on mitochondrial promoters being highly significant for each cell model (Fisher’s test, p value < 2.2e-16)(**Supplemental Fig. S2B)**. In accord with GPS2 preferential binding to promoters rather than enhancer and other inter/intragenic regions, GPS2 peaks on mitochondrial genes in 3T3-L1 cells showed a significant enrichment in promoter-associated motifs (**Supplemental Fig. S2C**). Also, tag density plots representing the overlapping of GPS2 ChIPseq profile with the positions of known nuclear partners of GPS2 (Cardamone et al., 2014; Fan and Evans, 2014; Hock and Kralli, 2009; Lefterova et al., 2010; Raghav et al., 2012) showed that the center of GPS2 binding was shifted towards the transcription start site (TSS) when compared to the binding of nuclear receptors and associated corepressors **(Fig. 2C**). This suggested that GPS2 recruitment on neMITO genes is distinct from what previously reported for other genomic targets (Cardamone et al., 2012; Cardamone et al., 2014; Fan et al., 2016). In keeping with this conclusion, comparative analysis of GPS2 binding with existing GROseq and H3K27Ac profiles in adipocytes (Harms et al., 2015; Step et al., 2014) confirmed that GPS2 binding was concentrated around the TSS of mitochondrial genes **(Fig. 2D)** where it overlapped with the binding profile of RNA Polymerase II (POL2)(Siersbaek et al., 2011)**(Fig. 2E)**. Collectively, integration of RNAseq and ChIPseq data therefore reveal that GPS2 regulates mitochondrial function through direct transcriptional regulation of a vast nuclear-encoded mitochondrial gene program.

### GPS2-mediated inhibition of Ubc13 activity is required for H3K9 demethylation and expression of neMITO genes

8

To elucidate the specificity of this regulatory strategy, we next asked what constituted the molecular mechanism of GPS2-mediated regulation of transcription in the context of neMITO genes. As GPS2 is a potent inhibitor of the ubiquitin conjugating enzyme Ubc13 and regulates the expression of selected PPARγ target promoters through stabilization of an H3K9 demethylase (Cardamone et al., 2012; Cardamone et al., 2014; Cederquist et al., 2017; Lentucci et al., 2017), we first investigated whether H3K9 demethylation and local ubiquitination are involved in the regulation of mitochondrial genes. This proved correct as in 3T3-L1 GPS2-KD cells we detected: i) a significant increase in the level of non-proteolytic ubiquitin chains on the promoters of mitochondrial genes *Ndufv1* and *Tomm20* by ChIP using an antibody specific for K63 ubiquitin chains **(Fig. 3A)**; ii) a dramatic increase in H3K9me3 occupancy **(Fig. 3B**). In agreement with the regulation observed on specific PPARγ-bound promoters (Cardamone et al., 2014), the accumulation of H3K9me3 on the promoters of *Ndufv1* and *Tomm20* correlated with a significant decrease in binding of the H3K9 demethylase JMJD2A/KDM4A (**Supplemental Fig. S3A**). However, our previous genome-wide data indicated that KDM4A binding is limited to a small subset of mitochondrial genes (Cardamone et al., 2014), whereas ChIPseq genome-wide analysis of H3K9 methylation status revealed that GPS2 downregulation caused a significant increase in H3K9me3 on the entire mitochondrial genes set **(Fig. 3C**). Notably, the increase in H3K9me3 in absence of GPS2 was specific to mitochondrial genes as revealed by comparison to the whole genome or to randomly selected group of genes of comparable size (**Fig. 3C** and **Supplemental Fig. S3B**). Also, intriguingly, the basal level of H3K9me3 in the area around neMITO genes was found significantly higher than for other genomic loci **(Fig. 3C** and **Supplemental Fig. S3B**). These results together indicate that neMITO genes in WT cells are specifically marked by the exclusion of H3K9me3 from the proximal promoter area, which is lost, or at least significantly reduced, upon GPS2 downregulation.

**Figure 3.**
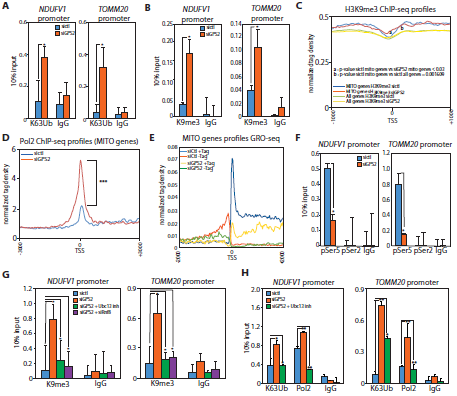
GPS2-mediated inhibition of K63 ubiquitination is required for H3K9 de-methylation, POL2 processivity and transcription of mitochondrial genes. **(A)(B)** Increased K63 ubiquitination (A) and H3K9me3 (B) in 3T3-L1 cells transfected with siGPS2 as measured by ChIP on the promoters of representative neMITO genes *Ndufv1* and *Tomm20*. **(C)**Tag density plot of H3K9me3 ChIPseq data showing a significant increase in H3K9me3 specific to mitochondrial genes (2086 genes) compared to the full genome (20950 genes in mm8 with length>1kb). P-values were computed using the Wilcoxon test based on the difference in normalized counts +/-200bp around the TSS. **(D)**Tag density plot of POL2 ChIPseq data showing a significant increase (p-value<2.99e-09) in POL2 binding on mitochondrial genes in 3T3-L1 cells transfected with siGPS2. **(E)**Tag density plot of GROseq data showing nascent transcription being impaired in GPS2-KD cells for both sense (+Tag) and anti-sense (-Tag) transcription **(F)**ChIP analysis of POL2 phosphorylation (Ser5 and Ser2) in 3T3-L1 cells transfected with siCTL or siGPS2. (G)Accumulation of H3K9me3 upon GPS2 downregulation is rescued either by treating cells with the Ubc13 inhibitor drug NSC697923 or by transient downregulation of ubiquitin E3 ligase RNF8. **(H)**Accumulation of K63 ubiquitin chains and POL2 binding upon GPS2 downregulation are rescued by treatment with the Ubc13 inhibitor drug NSC697923. ChIPseq data is representative of two independent biological replicates. Hand ChIPs data are representative of three independent experiments, bar graphs represent the sample mean of three technical replicates +/-SD. * indicate p-value<0.05 **indicate p-value<0.01, ***indicate p-value<0.001.

To our initial surprise, the accumulation of H3K9me3, which appeared concentrated downstream of the TSS, was associated with a significant increase in RNA Polymerase II (POL2) binding over mitochondrial promoters **(Fig. 3D and Supplemental Fig. S3C**). Notably, despite GPS2 binding to an extensive number of genes, a quantitatively significant increase in POL2 occupancy was observed only for the mitochondrial gene subset and not for comparable groups of random genes, just as observed for the H3K9me3 mark **(Supplemental Fig. S3D)**. This suggested that POL2 activity rather that its recruitment might be affected by the methylation status of H3K9.

As GPS2 was reported associated with the negative elongation factor NELF and paused RNA polymerase II (POL2) on HIV long terminal repeats (Natarajan et al., 2013), we asked whether the expression of mitochondrial genes could be impaired in GPS2-KD cells as a result of increased POL2 proximal-promoter pausing. To interrogate the status of POL2 on GPS2-regulated genes, we measured nascent RNA by GROseq in WT and GPS2-KD 3T3-L1 cells. In accord with the transcriptomic analysis, transcription of mitochondrial genes was found dramatically reduced upon GPS2 downregulation, with divergent polymerase molecules driving sense and anti-sense transcription being equally impaired (**Fig. 3E** and **Supplemental Table 3**). However, the read density pattern of GROseq tags observed in GPS2-KD cells resembled what previously reported in cells treated with the initiation inhibitor Triptolide, rather than the pattern reported upon treatment with the pausing escape inhibitor Flavopiridol (Chen et al., 2015; Jonkers et al., 2014). Thus, we conclude that the lack of significant accumulation of nascent RNA detected over the first 50-100bp of mitochondrial genes indicates that GPS2 downregulation does not promote an increase in POL2 pausing, but rather inhibits POL2 activation **(Fig. 3E)**. In accord with this conclusion, profiling the phosphorylation status of Ser2 and Ser5 sites on the C-terminal domain (CTD) of POL2 by ChIP confirmed that neither POL2 phosphorylation event was detectable on representative mitochondrial gene promoters in GPS2-KD cells, despite the increase in POL2 accumulation (**Fig. 3F**). Notably, the transcription of mitochondrial genes up-regulated in GPS2-KD cells by RNAseq was also found impaired by GROseq (**Supplemental Fig. S3E**). This result confirms our initial hypothesis that GPS2 is globally required for the transcriptional activation of neMITO genes and suggests that the increased mRNA levels observed for a subset of mitochondrial genes by RNAseq profiling result from a secondary response, possibly involving post-transcriptional mechanisms of RNA stabilization. In conclusion, integration of our ChIPseq and GROseq experiments reveals that GPS2 depletion causes a robust accumulation on the promoters of neMITO genes of a hypo-phosphorylated and unproductive form of POL2, which is unable to escape from the core promoter and promote productive transcription, likely because of the altered chromatin environment aberrantly enriched in the repressive histone mark H3K9me3.

Lastly, we investigated whether GPS2 regulates chromatin remodeling and POL2 activation through modulation of Ubc13-mediated ubiquitination. In support of this hypothesis, inhibition of Ubc13 enzymatic activity in GPS2-KD cells with the specific inhibitor NSC697923 (Liao et al., 2015; Mallipattu and He, 2015; Pulvino et al., 2012; Small et al., 2011), or downregulation of the ubiquitin ligase RNF8 via siRNA (Cardamone et al., 2014), were sufficient for rescuing the aberrant accumulation of H3K9me3, K63 ubiquitin chains and the stalling of POL2 on target mitochondrial promoters (**Fig. 3G** and **3H**). Thus, our results collectively support the existence of a regulatory strategy, specific for neMITO genes, which is based on GPS2 being required for preventing the accumulation of the repressive mark H3K9me3 and the stalling of POL2, via inhibition of Ubc13-mediated synthesis of K63 ubiquitin chains.

### Retrograde translocation of GPS2 upon mitochondria depolarization

These results establish that GPS2 is required to sustain the synthesis of nuclear-encoded mitochondrial components through its essential role in promoting gene expression of mitochondrial proteins encoded in the nucleus. However, GPS2 does not appear to be required for the expression of genes encoded in the mitochondrial DNA, thus raising the question of what is the function of GPS2 physical presence in the mitochondria. As the expression of neMITO genes has to be regulated in response to the functional needs of mitochondria, we became intrigued by the possibility that mitochondrial GPS2 might play a role in integrating signals from the mitochondria with the regulation of nuclear transcription, a function known as mitochondrial retrograde signaling (Guha and Avadhani, 2013; Jazwinski, 2012; Whelan and Zuckerbraun, 2013). To explore the hypothesis that GPS2 functions as a mediator of mitochondria retrograde signaling, we first used the uncoupling agent carbonilcyanide p-triflourometho-xyphenylhydrazone (FCCP) to depolarize the mitochondria. Mild treatment with FCCP induces mitochondria-to-nucleus signaling to promote the activation of an integrated stress response program that include the transcriptional activation of nuclear-encoded mitochondrial genes as needed for sustaining the metabolic reprogramming of depolarized cells and the biogenesis of new mitochondria (Catic et al., 2013; Quiros et al., 2017; Yun and Finkel, 2014). After 3 hours treatment with FCCP, we observed a significant decrease in GPS2 localization in the mitochondria, as shown by loss of co-staining with mtHSP70, and a corresponding increase in nuclear staining for GPS2 (**Fig. 4A**). As expected based on known GPS2 association with nuclear receptors and corepressors transcriptional functions (Cardamone et al., 2012; Cardamone et al., 2014; Guo et al., 2014; Toubal et al., 2013), non-stressed cells already presented a significant amount of basal nuclear staining (**Fig. 4A**). Thus, to best appreciate the increase in nuclear GPS2, we examined GPS2 localization through subcellular fractionation and western blotting. While total GPS2 protein levels did not change upon mitochondria depolarization (**Supplemental Fig. S4A**), blotting of fractionated extracts confirmed a significant redistribution of GPS2 from the mitochondria to the nucleus (**Fig. 4B**).

**Figure 4.**
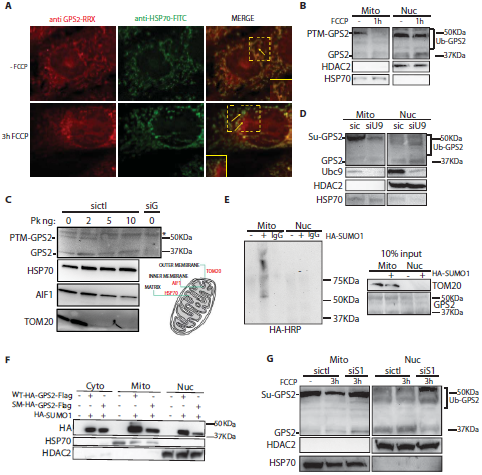
Mitochondria-to-nucleus regulated translocation of GPS2 in response to mitochondria depolarization. **(A)**Co-localization of GPS2 and mtHSP70 by IHF staining of 3T3-L1 cells shows reduced GPS2 localization to the mitochondria and corresponding increased nuclear staining (see inset and arrows) upon 3h of FCCP. **(B)**Translocation from mitochondria to nucleus as shown by western blotting of fractionated extracts in wild type 3T3-L1 cells treated with FCCP for 1h. Distinct post-translationally modified band are indicated on the side of the blot. WBs for HDAC2 and mtHSP70 respectively serve as control for nuclear (Nuc) and mitochondrial (Mito) extracts. **(C)**Proteinase K (Pk) assay showing localization of GPS2 and post translational modified GPS2 (PTM-GPS2) in the mitochondria matrix and outer membrane respectively. The degradation patterns of TOM20, AIF1 and HSP70 are shown as respectively markers of the OMM, the IMM and the mitochondrial matrix compartments. **(D)**GPS2 post-translational modification in mitochondria is reduced with downregulation of the SUMO E2 enzyme Ubc9 (siU9). **(E)***In vivo* Sumoylation assay in 293T cells showing specific sumoylation of GPS2 with HA-SUMO1 in mitochondrial extrcts. **(F)**Mitochondria localization of overexpressed wild type HA-GPS2-Flag and the HA-GPS2-Flag K45,K71R Sumo mutant (SM) by WB of fractionated extracts in 293T cells. **(G)**Impaired translocation of GPS2 in depolarized-treated cells transfected with siSENP1 (siS1) to block the desumoylation of GPS2.

Unexpectedly, this approach revealed that GPS2 id differentially modified in the various cellular compartments, with the specificity of each band being confirmed by GPS2 downregulation through two independent siRNAs **(Supplemental Fig. S4B**). In particular, most of the endogenous mitochondrial GPS2 (mtGPS2) was found to run significantly higher than the expected MW of 36kD, which indicated that mtGPS2 might be specifically marked through a specific post-translational modification, likely distinct from those observed in the nucleus (**Fig. 4B** and **Supplemental Fig. S4B**). Furthermore, comparing the pattern of degradation observed for mtGPS2 and known markers of the different compartments when subjecting isolated mitochondria to digestion with increasing amounts of proteinase K (PK) actually revealed the existence of two pools of mtGPS2 (**Fig. 4C**). One pool one was localized to the matrix and consisted of unmodified GPS2. The other pool was localized to the outer mitochondrial membrane (OMM) and consisted of the post-translationally modified GPS2 (PTM-GPS2)**(Fig. 4C**). Localization of these two pools was further confirmed by differential trypsin digestions sensitivity of isolated mitochondria (Mt) or mitoplast (MP) lacking the OMM **(Supplemental Fig. S4C**).

### Regulation of GPS2 intracellular localization through sumoylation/desumoylation

Because the intracellular re-localization of GPS2 observed upon depolarization involved a decrease of PTM-GPS2 in the mitochondria associated with an increase in unmodified GPS2 in the nucleus (see **Fig. 4B**), we reasoned that mitochondria-to-nuclear translocation of GPS2 could be modulated by post-translational modifications. Based on the size of the shift, on previous reports of GPS2 being sumoylated on residues K45/K71(Bi et al., 2014; Huang et al., 2015), and consistent with the important role played by SUMO enzymes in the regulation of mitochondrial function and dynamics (Braschi et al., 2009; Fu et al., 2014; Prudent et al., 2015; Zunino et al., 2007), we asked whether GPS2 sumoylation accounts for the PTM observed in the mitochondria. This hypothesis was confirmed by the following experiments: i) downregulation of the sumo-conjugating enzyme UBC9 was sufficient to observe a significant decrease in the amount of PTM-GPS2 present in mitochondrial extracts (**Fig. 4D**); ii) sumoylation of endogenous GPS2 in the mitochondrial fraction was observed in IP/sumoylation assay with HA-SUMO1 **(Fig. 4E**), iii) sumoylation of the overexpressed GPS2, in presence of HA-SUMO1, was abrogated by specific mutation of K45 and K71 (K45,71R mutant (Huang et al., 2015), **Fig. 4F**). Together, these results indicate that sumoylation on K45/K71 accounts for the shift in GPS2 size in mitochondria. In addition, as downregulation of UBC9 did not affect GPS2 higher molecular weight bands in the nucleus (**Fig. 4D**), our results indicate that sumoylation is restricted to the mitochondria (see also **Fig.4E**), whereas nuclear GPS2 is modified by other means, including ubiquitination as previously reported (Huang et al., 2015). On the contrary, there was a small but consistent increase in nuclear unmodified GPS2 upon UBC9 downregulation (**Fig. 4D**). This observation suggested that GPS2 sumoylation status could be important for determining its intracellular localization, with sumoylation promoting GPS2 localization out of the nucleus. This hypothesis was supported by the decrease in SUMO-GPS2 observed upon FCCP-induced relocalization of GPS2 (see **Fig. 4B**). Thus, we asked whether GPS2 desumoylation was required for its nuclear translocation upon FCCP treatment. Indeed, downregulation of the SUMO protease SENP1, which was previously reported to desumoylate GPS2 (Bi et al., 2014), was sufficient to promote not only the stabilization of SUMO-GPS2, but also its retention in the mitochondria (**Supplemental Fig. S4D** and **Fig. 4G**), confirming the role of sumoylation in regulating GPS2 subcellular localization.

### GPS2 is required for the transcriptional activation of a stress response nuclear program upon mitochondria depolarization

These results together confirm that, upon loss of mitochondrial membrane potential, GPS2 translocates from mitochondria to nucleus in a desumoylation regulated fashion. We next sought to investigate whether GPS2 relocalization is important for regulating the nuclear transcription response to mitochondrial stress. Recently, integrative analysis of Hela cell proteomic, transcriptomic and metabolic profiling has defined a comprehensive stress response activated in response to long term exposure to a number of mitochondrial stressors, including FCCP (Quiros et al., 2017). To discriminate between primary and secondary events within this response and assess to which extent GPS2 retrograde translocation is required for the activation of a nuclear transcriptional program, we analyzed the early transcriptional events induced upon FCCP treatment by GROseq in WT and GPS2-KD 3T3-L1 cells. Using an FDR of 5%, we identified 1486 upregulated genes and 1060 downregulated genes upon 3 hours of FCCP treatment, (**Fig. 5A** and **Supplemental Table 3**). Mitochondrial genes were similarly represented among both up-and down-regulated groups (205 upregulated mitochondrial genes and 163 downregulated mitochondrial genes). In agreement with the published work, FCCP stimulation induced the activation of genes associated with “Folate metabolism”, “tRNA aminoacylation”, “Aminoacid metabolism”, “Carbon metabolism” and “Ubiquitin-mediated proteolysis” (**Fig. 5A**, **Supplemental Fig. S5A** and **Supplemental Table 3**)(Quiros et al., 2017). In comparison, downregulated genes instead associated with “Cell cycle”, “Cholesterol and Lipid Metabolism”, “Glucose metabolism” and “Apoptosis” (**Fig. 5A**, **Supplemental Fig. S5A** and **Supplemental Table 3**). Among downregulated mitochondrial genes we also observed a number of genes associated with “Oxidative phosphorylation” and “Mitochondrial Ribosomal Proteins”, categories found decreased by proteomic analysis at later times (Quiros et al., 2017), however the enrichment did not reach statistical significance. Together these results indicate that the early response to mitochondrial depolarization promotes cellular reprogramming through the regulation of a vast transcriptional program.

**Figure 5.**
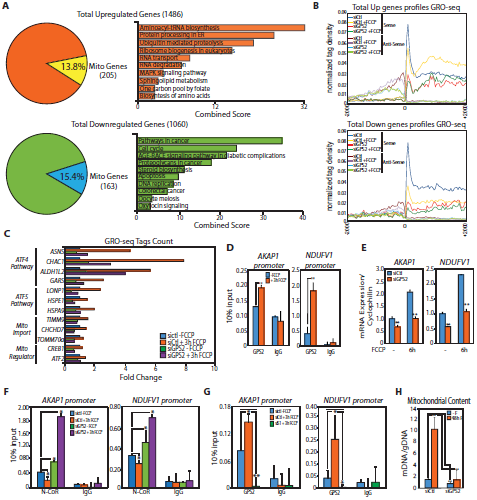
Impaired nuclear response to mitochondrial stress upon GPS2 depletion. **(A)**DEGs defined by GROseq between untreated and FCCP-treated 3T3-L1 cells (3h), including analysis of 10 most enriched pathways among upregulated and downregulated genes (EnrichR, KEGG Pathways). **(B)**Tag density plot of GROseq data showing transcriptional activation of DE upregulated genes upon FCCP is abrogated in GPS2-KD cells. Transcription of FCCP-downregulated genes is already reduced in GPS2-KD cells. **(C)**Activation of representative genes of the ATF4-mediated integrated stress respose, ATF5-mediated mitochondrial unfolded protein response, mitochondria import machinery and regulators of mitochondria retrograde signaling depends on GPS2 as measured by GROseq. **(D)**ChIP analysis of GPS2 recruitment to mitochondrial genes and NCoR targets *AKAP1* and *NDUFV1* upon 3h of FCCP. **(E)**Relative expression of mitochondrial genes and NCoR targets *AKAP1* and *NDUFV1*upon 6h of FCCP as measured by RT-qPCR in WT and GPS2-KD3T3-L1 cells. **(F)**Dismissal of corepressor NCoR from target genes *AKAP1* and *NDUFV1* upon 3h FCCP treatment is impaired upon GPS2 downregulation by ChIP. **(G)**Increased GPS2 binding to nuclear-encoded mitochondrial genes upon 3 hours of FCCP stimulation is impaired in cells transfected with siSENP1 (siS1). **(H)**Increased in mitochondrial DNA content upon long term FCCP treatment (48h) is impaired in GPS2-KD 3T3-L1 cells as measured by PCR (ratio of mtDNA/nDNA). ChIP results in **(D), (F)** and **(G)** are representative of three independent experiments, bar graphs represent the sample mean of three technical replicates +/-SD. Results in **(E)** and **(H)** represent the sample mean of three independent experiments; error bars represents.e.m. (n=3). * indicate p-value<0.05; ** indicate p-value<0.01.

Comparing this signature between WT and GPS2-KD cells revealed that the basal transcription of both upregulated and downregulated genes, as well as the activation of the upregulated genes upon FCCP required GPS2 (**Fig. 5B**). This indicates that GPS2 translocation to the nucleus was broadly required for supporting the transcriptional response to mitochondrial stress. In fact, the list of DEGs upon 3h of FCCP included known targets of both the ATF4-dependent integrated stress response and ATF5-dependent mtUPR response (Fiorese et al., 2016; Nargund et al., 2012; Quiros et al., 2016; Quiros et al., 2017), in addition to proteins important for mitochondrial protein import and TFs previously associated with retrograde signaling and mitochondria biogenesis (Arnould et al., 2015; Biswas et al., 1999; Than et al., 2011; Vankoningsloo et al., 2006)(**Fig. 5C**). Because the activation of genes within all these groups was either completely lost or significantly impaired in GPS2-KD cells, we checked GPS2 binding to their promoters and found peaks on each promoter, in agreement with GPS2 playing a broad role in the transcriptional regulation of a nuclear stress response to mitochondrial depolarization by FCCP in mammalian cells (**Supplemental Fig. S5B**).

To our initial surprise, the GROseq did not reveal any significant upregulation of NCoR/CREB mitochondrial target genes previously reported as activated in response to FCCP through regulation of NCoR stability (Catic et al., 2013). However, further characterization by ChIP following FCCP treatment confirmed a significant increase in GPS2 binding to representative promoters of *AKAP1* and *Ndufv1* neMITO genes after 3h of FCCP treatment (**Fig. 5D)**. This was followed by a GPS2-dependent, corresponding increase in gene expression at later time points (**Fig. 5E**). Also, ChIP analysis confirmed that FCCP-induced activation of both *AKAP1* and *Ndufv1* genes associated with the dismissal of NCoR, whereas in GPS2-KD cells repressed genes were marked by increased NCoR binding (**Fig. 5F**). Most importantly, the upregulation of these neMITO genes upon depolarization depended on GPS2 retrograde translocation from the mitochondria as GPS2 recruitment to the promoters of *NDufv1* and *AKAP1* genes was blocked by downregulation of the SUMO protease SENP1 (**Fig. 5G**). These results, together, confirm that GPS2 is a mediator of mitochondria retrograde signaling and reveal that its regulated translocation from mitochondria to the nucleus is required for promoting the transcriptional activation of an extensive transcriptional program that includes both stress response genes as well as mitochondrial genes, as required for responding to the mitochondrial damage through cellular reprogramming and mitochondria biogenesis. Accordingly, the increase in mitochondrial content induced during the recovery response to FCCP depolarization was dramatically impaired in GPS2-KD cells (**Fig. 5H**).

### Retrograde translocation of GPS2 during adipogenesis

Based on these results, we reasoned that mitochondria-to-nucleus translocation of GPS2 might not be limited to stress-response events. To explore the role of GPS2 in the activation of mitochondrial gene expression under physiological conditions, we investigated GPS2 nuclear function and retrograde translocation during 3T3-L1 adipogenic differentiation, a process known to elicit significant mitochondrial biogenesis (Raghav et al., 2012; Wilson-Fritch et al., 2003; Yu et al., 2005). Our prediction was that GPS2 would translocate from mitochondria to the nucleus during the differentiation of 3T3-L1 cells in order to support mitochondria biogenesis through transcriptional activation of the mitochondrial gene program. Consistent with this prediction, we observed: i) a 50% increase in GPS2 binding to the promoters of mitochondrial genes when comparing GPS2 ChIPseq data from 3T3-L1 cells at day0 and day6 of differentiation (Cardamone et al., 2014)(**Fig. 6A**); ii) a significant reduction in the activation of a number of adipo-specific mitochondrial genes in GPS2-depleted, PPARg-expressing pre-adipocytes (**Fig. 6B**). Accordingly, the increase in mitochondrial content normally observed during the early phase of differentiation was almost completely abrogated upon GPS2 downregulated (**Fig. 6C**).

**Figure 6.**
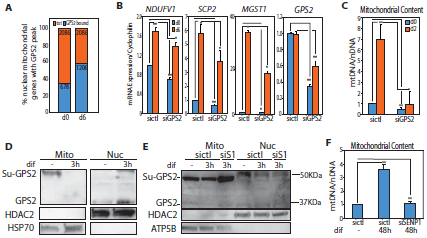
GPS2 retrograde translocation supports mitochondrial biogenesis during adipogenesis. **(A)**GPS2 binding to nuclear-encoded mitochondrial genes as measured by ChIPseq increases upon differentiation (GSE57779). **(B)**RT-qPCR analysis of the expression of adipo-specific mitochondrial genes in differentiating 3T3-L1 adipocytes (day0 and day6 of differentiation). **(C)**Decreased mitochondrial biogenesis during 3T3-L1 adipogenesis in GPS2-KD cells as measured by increase in mitochondrial DNA content (ratio of mtDNA versus genomic DNA). **(D)**GPS2 translocation from mitochondria to nucleus as measured by western blotting in subcellular fractionated extracts 3h after induced adipogenesis. **(E)(F)** GPS2 desumoylation and translocation from mitochondria to nucleus (E) and mitochondrial biogenesis (F) during adipogenesis is impaired in cells transfected with siSENP1 (siS1). Results in **(A), (B)** and **(F)** represent the sample mean of three independent experiments; error bars represent s.e.m. (n=3). *indicate p-value<0.05; **indicate p-value<0.01.

As observed in the case of FCCP-treated cells, GPS2 engagement for the expression of neMITO genes was associated with a rapid decrease in SUMO-modified GPS2 in the mitochondria parallel to a corresponding increase in unmodified GPS2 in the nucleus shortly after induction of differentiation with a classic adipogenic cocktail (**Fig. 6D**). Furthermore, in keeping with the hypothesis that desumoylation is required for the nuclear translocation of mitochondrial GPS2, the downregulation of the desumoylation enzyme SENP1 was sufficient to inhibit both the translocation from mitochondria to nucleus and the resulting increase in mitochondria content (**Fig. 6E and 6F**). Therefore, we conclude that GPS2 shuttling between mitochondria and nucleus is a regulatory strategy important for promoting the nuclear expression of mitochondrial genes not only in response to mitochondrial stress, but also during the physiological metabolic reprogramming associated with cell differentiation, in this case of adipocytes.

### Reduced mitochondrial content in the BAT of GPS2-AKO mice

Finally, to confirm the *in vivo* relevance of this regulatory strategy, we analyzed the brown adipose tissue (BAT) of adipo-specific GPS2-AKO mice. Adipose tissue-specific GPS2 deletion promotes obesity associated with constitutive insulin signaling, increased lipid deposition in the white adipose tissue (WAT) and improved systemic insulin sensitivity (Cederquist et al., 2017). In agreement with the altered ratio of lipogenesis/lipolysis observed in the WAT, inspection of the BAT revealed that GPS2 deletion promoted whitening of the brown depot (**Fig. 7A**). A significant increase in the size of lipid droplets was also observed in brown adipocytes differentiated *in vitro* from the stroma vascular fraction (SVF) of GPS2-AKO mice compared with WT littermates (**Fig. 7B**). To investigate whether this phenotype reflected a defect in mitochondria biogenesis, we measured mtDNA copy number in BAT from wild type and GPS2-AKO mice, and found that GPS2-deficient mice had only 50% mtDNA levels compared to their WT littermates (**Fig. 7C**). However, respiration measured in isolated mitochondria from the BAT tissue was not impaired in absence of GPS2 (**Fig. 7D**). These results indicate that mitochondria functionality is not impaired in GPS2-deficient BAT, but rather there is a decrease in mitochondrial mass, which was confirmed by measuring the amount of mitochondrial proteins in the tissue (**Fig. 7E and 7F**). Also in agreement with this conclusion and with our *in vitro* studies, we observed a significant downregulation of neMITO genes in the BAT tissue (**Fig. 7G**). Mechanistically, ChIPseqs analysis of H3K9me3 performed on the BAT from WT and GPS2-AKO littermates confirmed a significant increase in H3K9me3 on the promoters of neMITO genes in the GPS2-deficient depot (**Fig. 7H**). Furthermore, the increase in H3K9 trimethylation correlated with aberrant accumulation of POL2, as measured by ChIP on the promoters of mitochondrial genes *Ndufv1* and *Tomm20* (**Fig. 7I**), thus fully confirming *in vivo* the molecular mechanism described *in vitro* in 3T3-L1 cells.

**Figure 7.**
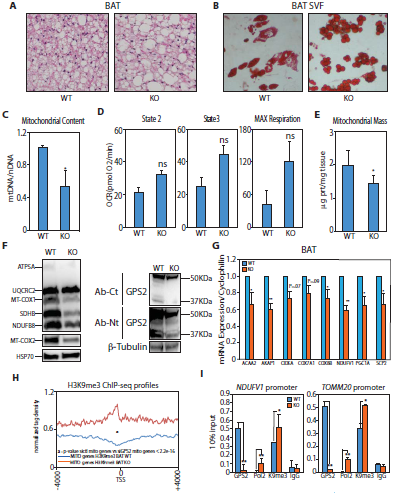
GPS2 regulates mitochondrial biogenesis *in vivo* in Brown Adipose Tissue. **(A)**H&E of BAT from WT and KO littermates mice showing increased lipid deposition in the adipose tissue of GPS2-AKO. **(B)**Oil Red O staining of *in vitro* differentiated adipocytes using SVF isolated from the brown adipose tissue of WT and KO littermates. **(C)**Decreased mitochondrial DNA content in the BAT of GPS2-AKO mice compared to WT littermates. **(D)**Quantification of oxygen consumption rates (OCR) in BAT isolated mitochondria from chow-fed WT and GPS2-AKO littermates under the different respiratory states. State 2 quantifies respiration driven by proton leak, state 3 quantifies respiration linked to maximal ATP synthesis and MAX respiration represents maximal electron transport chain activity induced by FCCP. Results represent average± SEM for complex I-driven respiration (pyruvate-malate, n=6 mice per group), complex II-driven respiration (succinate-rotenone, n=6 mice per group). **(E)**Reduced mitochondrial protein content in BAT isolated from GPS2-AKO compared to WT littermates mice. **(F)**Western blot analysis of mitochondrial protein expression in BAT from GPS2-AKO compared to WT mice. Western Blot using two different antibodies against GPS2 is shown as a control of GPS2 deletion in GPS2-AKO mice. The deletion, driven by Adipoq-Cre, is specific to the mature adipocytes not other cell types in the tissue. **(G)**RT-qPCR analysis of representative nuclear-encoded mitochondrial genes in the BAT of WT and KO mice. **(H)**Tag density plot of H3K9me3 ChIPseq performed on BAT from WT and KO littermates showing a significant increase in H3K9me3 on the promoters of mitochondrial genes in GPS2-AKO mice. **(I)**ChIP for H3K9me3, POL2 and GPS2 showing increased promoter occupancy by POL2 and H3K9me3 in GPS2-AKO BAT on representative genes *Ndufv1* and *Tomm20*. Results in **(C), (E)** and **(G)** represent the sample mean of five independent experiments; error bars represent s.e.m. (n=5). ChIPs data are representative of four independent experiments, bar graphs represent the sample mean of three technical replicates +/-SD. *indicate p-value<0.05; **indicate p-value<0.01.

## Discussion

Mitochondrial mass and functions differ considerably among tissues and are dynamically regulated in response to different physiological cues, including nutrient availability, cold exposure and endurance exercise (De Pauw et al., 2009; Hock and Kralli, 2009). Mitochondrial biogenesis and remodeling are also critical for embryonic and adult stem cells differentiation (Wanet et al., 2015). Because the majority of mitochondrial proteins are encoded in the nuclear genome, the assembly of new mitochondria and the regulation of the metabolic functions of existing mitochondria critically depends on the transcriptional regulation of mitochondrial gene expression in the nucleus (anterograde signaling) and feedback signaling from the mitochondria to the nucleus (retrograde signaling)(Finley and Haigis, 2009; Jazwinski, 2013; Kotiadis et al., 2014; Whelan and Zuckerbraun, 2013). A major conclusion of this work is that it adds GPS2 to the existing network of transcription factors and cofactors regulating the expression of mitochondrial gene expression in the nucleus. Our results in fact link the nuclear actions of GPS2, and its ability to inhibit the K63 ubiquitination machinery, to chromatin remodeling events controlling the transcriptional activation of neMITO genes by RNA POL2. Notably, the fact that GPS2-mediated regulation occurs at the level of nuclear core promoters, adds a novel layer to the known mechanisms regulating the transcriptional control of mitochondrial genes. Indeed, our data suggest that the complexity and specificity of mitochondrial gene expression is achieved through multiple levels of regulation mediated not only by distal regulators of transcription (i.e. nuclear receptors and associated cofactors like PGC1α and PGC1β(Fan and Evans, 2014; Hock and Kralli, 2009; Lin et al., 2005)), but also by proximal regulators of transcription, such as GPS2. Future studies will be required to address how these layers of regulation are coordinated to maintain cellular homeostasis.

The current study also indicates that neMITO genes are specifically sensitive to the methylation status of H3K9, revealing an unexpected role for H3K9me3 in the transcriptional regulation of active genes in addition to the well known roles in gene silencing and heterochromatin organization. As histone demethylases JMJD-1.2 and JMJD-3.1 have been recently described as key mediators of lifespan extension and mtUPR response in worms (Merkwirth et al., 2016), these results suggest that mitochondrial gene expression is specifically sensitive to a conserved regulatory strategy based on the modulation of H3K9 histone methylation/demethylation. Our data indicate that in mammalian cells, demethylation is partially achieved, but not limited to the action of the histone demethylase JMJD2/KDM4A. Other demethylases that are known to play a role in the regulation of mitochondrial functions are likely to contribute (Duteil et al., 2016; Hino et al., 2012; Liu and Secombe, 2015; Merkwirth et al., 2016; Tian et al., 2016). In fact, the use of specific demethylases might be important for providing specificity to the regulation of different subsets of mitochondrial genes.

Another fundamental finding of this study is that GPS2 fulfills the role of a direct mediator of mitochondrial retrograde signaling in mammalian cells. Its transcriptional activity is in fact directly regulated and informed by the mitochondria functional status through regulated mitochondria-to-nucleus translocation. The mitochondria retrograde signaling pathway was initially identified in *S. Cerevisiae* as an adaptation mechanism that allows yeast cells to respond to mitochondrial dysfunctions by activating a transcriptional program that includes both metabolic and stress response genes (Bohovych and Khalimonchuk, 2016; Guha and Avadhani, 2013; Jazwinski and Kriete, 2012; Whelan and Zuckerbraun, 2013). In yeast, the retrograde factors responsible for signaling through the retrograde pathway are the DNA-binding transcription factors Rtg1-3 and a regulatory subunit, Rtg2, which senses changes in mitochondrial membrane potential, regulates the nuclear import of Rtg1-3 and directly contributes to the transcriptional regulation of the retrograde gene program in the nucleus by participating in the SLIK chromatin remodeling complex (Jazwinski, 2005; Jazwinski and Kriete, 2012; Liao and Butow, 1993; Miceli et al., 2011; Sekito et al., 2000). While the extensive search for homologs of the yeast retrograde genes in higher organism has revealed possible overlapping functions of Rtg1/Rtg3 with mammalian transcription factors, no homolog has thus far been identified for Rtg2 (Jazwinski, 2013). Based on the results reported here, we propose that GPS2 act as a functional homolog of Rtg2 in mammalian cells. As with Rtg2, GPS2 represents a direct link between the nuclear transcription factors regulating mitochondrial gene expression and the mitochondria itself. In a similar manner to Rtg2 regulation, GPS2 function and nuclear translocation is triggered by mitochondria depolarization. Also, in accordance with Rtg2 contribution to the chromatin remodeling related to the yeast SLIK (SAGA-like) complex, GPS2 directly binds to and modulates the chromatin environment of target gene promoters. Intriguingly, it does so by preventing the accumulation of H3K9me3, which could be inhibitory against H3K9ac, a mark of active transcription deposited by Gcn5/PCAF, the catalytic subunit of the SAGA complex. In addition, Rtg2-mediated regulation of Rtg1-3 nuclear import via modulation of an inhibitory phosphorylation cascade closely resembles GPS2-mediated regulation of NFkB activity, as well as our recent finding of GPS2 promoting FOXO1 stabilization and its nuclear functions via inhibition of AKT ubiquitination and activation (Cardamone et al., 2012; Cederquist et al., 2017; Lentucci et al., 2016). Lastly, our *in vivo* data indicate that GPS2 plays a critical role in the regulation of lipid metabolism and energy expenditure in the adipose tissue, just as RTG2 has proven to be an essential factor for the regulation of nutrient metabolism in yeast cells. In both cases, the phenotypic output is achieved through the coordinated regulation of an extensive transcriptional program, including genes involved in mitochondrial biogenesis, lipid metabolism and possibly other functions required for the metabolic reprogramming related to changes in mitochondrial function.

While these parallelisms are striking, they also raise numerous additional questions, including what is the nature of the relationship between GPS2 and the SAGA complex on nuclear target genes? What is the identity of GPS2 interacting partners in the mitochondria? And, how is sumoylation, possibly together with other regulatory strategies, modulating GPS2 tethering to the mitochondria and its shuttling between nucleus and mitochondria? In addition, future studies will be required for dissecting GPS2 role in integrating multiple pathways regulating cell growth, inflammation, metabolism, and stress resistance through regulation of ubiquitin signaling. This is going to be particularly relevant in the context of the adipose tissue as GPS2 was found significantly downregulated in human adipocytes from obese patients (Toubal et al., 2013). The loss of GPS2-mediated regulation of gene repression is in fact currently thought to contribute to the chronic inflammatory status of the obese adipose tissue. However, the studies reported here, together with our recent characterization of GPS2 role in regulating insulin signaling (Cederquist et al., 2017), add to the significance of GPS2 in the adipose tissue suggesting that it might serves a more general function in integrating the regulation of lipids mobilization from TG storage with their utilization via mitochondrial oxidative metabolism. As therapeutic and lifestyle interventions designed for improving insulin sensitivity often rely on increased mitochondria biogenesis and oxidative metabolism (Hu and Liu, 2011; Kusminski and Scherer, 2012; Newsholme et al., 2012; Patti and Corvera, 2010; Zamora et al., 2015), unraveling the molecular mechanisms regulating mitochondria biogenesis and function in response to energetic and metabolic demands is not only important for our basic understanding of cell biological processes, but also have the potential of leading to the development of novel therapeutic approaches for obese and diabetic patients.

## Authors Contribution

Conceptualization, V.P. and M.D.C.; Investigation, M.D.C, C.T.C, J.H., and K.M.; Formal Analysis, B.T.; Methodology, W.L.; Writing – Original Draft, V.P. and M.D.C.; Writing – Review & Editing, V.P., M.L. and M.G.R; Resources, M.G.R; Supervision,V.P. and M.L.

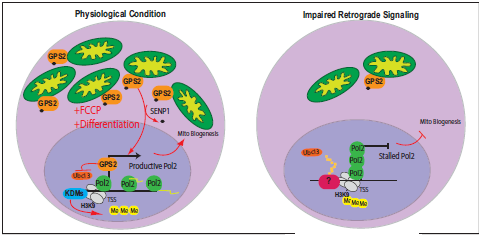

## Acknowledgments

We are extremely grateful to our colleagues in the Departments of Biochemistry and Medicine at Boston University and to past and present members of the Perissi lab for shared reagents and insightful discussions. We are also grateful to Dr. M. Picard (Columbia University) for reading the manuscript and providing valuable comments. We thank K. Ohgi and the UCSD sequencing core for assistance in preparing the libraries and running the next-generation sequencing experiments, the Analytical Instrumentation Core at Boston University for assistance in setting up experiments on the XF24 Seahorse Analyzer, the Confocal Microscopy Core at Boston University and Electron Microscopy Core at Harvard University for assistance with imaging experiments and Dr. G. Gaietta (UCSD) for initial guidance on imaging assays. This work was supported by Pilot and Feasibility Awards from Joslin/Boston University (V.P.) and from the Boston Nutrition and Obesity Research Center (BNORC)(V.P. and M.D.C.), by NRSA Individual Predoctoral Fellowship F31DK108571 (C.T.C.) and by NIH/NIDDK Research Grant R01DK100422 (V.P.)

## METHODS

### Reagents, Antibodies and siRNA

Anti-GPS2 antibodies were generated in rabbit against specific peptides representing aa 307-327 (Ct) and aa 1-11 (Nt). Representative ChIPs and IFs shown in the paper have been done using the Ct antibody, WBs using the Nt antibody, however it should be noted that all type of experiments have been carried out in the laboratory using either antibody with no difference noticed. Anti-N-CoR previously described^1,2^. The following antibodies were obtained from Santa Cruz Biotechnology: anti-HDAC2 (H54), TOM20 (F-10) and Ucb9 (N-19). Additionally, HSP70 (JG1-Thermo Scientific), beta-Tubulin (Sigma), ATP5B (A21351-Molecular Probes), Pol-II (ac-055-100-adgenode) K63-linkage-specific (HWA4C4-Enzo Life Science), KMD4A-ab47984, OXPHOs mix-ab110413, Pol-II phospho-ser5-ab5131, Pol-II phospho-ser2-ab24758 (Abcam), H3K93me (6F12-H4-Enzo Life Science) and mouse anti-HA (Upstate Biotechnology) were used in immunoblotting and immunostaining experiments. The siRNAs specific for GPS2, RNF8 and SENP1 were purchased from Invitrogen (*Silencer* ^*®*^ Select siRNAs), siRNA specific for GPS2 and RNF8 had been previously validated^3^. 2μM Ubc13 inhibitor NSC697923 (Sigma) and 10μM or 25μM Carbonyl cyanide-*p*-trifluoromethoxyphenylhydrazone(FCCP-TOCRIS) was used for cell treatments. Oligomycin A and Antimycin were purchased from Sigma.

### Animals studies

Fat-specific GPS2 knockout mice (GPS2-AKO) were generated using a cre/lox approach and maintained on a mixed 129sv/C57BL6J background^4^. Briefly, conditional Gps2 floxed mice were generated by inGenious Targeting Laboratory and adipose tissue specific deletion was achieved by crossing Gps2^flox/flox^ mice with heterozygous Adipoq-Cre C57BL/6J transgenic mice expressing Cre recombinase under control of the adiponectin promoter^5^. Male mice and littermate controls were used for all experiments. Mice were maintained on standard laboratory chow diet in temperature-controlled facility on a 12-hour light/dark cycle. All animal studies were approved by the Boston University Institutional Animal Care and Use Committee (IACUC) and performed in strict accordance of NIH guidelines for animal care.

### H&E staining

Upon harvesting, brown adipose tissue was incubated at 4˚C in Z-fix solution (Anatech LTD) overnight. Tissues were then transferred to 70% ethanol, paraffin embedded, sectioned, and stained with hematoxylin & eosin (Tuft Pathology Core/BNORC Adipose Biology Core). Imaging was performed as previously described^4^. **Cell culture and cell stainings.** Hela cell and 293T cell were grown in 10% FBS/DMEM supplemented with 0.1 mM MEM non-essential aminoacids, 2mM L-glutamine, 1mM sodium pyruvate. 3T3-L1 cells were grown in 10% BCS/DMEM supplemented with 0.1 mM MEM non-essential aminoacids, 2mM L-glutamine, 1mM sodium pyruvate. For cells transfection, Lipofectamine 2000 was used following the manufacturer’s protocol (Invitrogen). Induction of adipogenesis was performed in 10% FBS/DMEM high glucose supplemented with 0.1 mM MEM non-essential aminoacids, 2mM L-glutamine, 1mM sodium pyruvate, 0.1% 1mg/ml insulin, 0.01% 1mg/ml dexamethasone, 0.2% 55,6mg/ml isobutylmethylxanthine. Immunostaining was performed following standard protocols on cells fixed in 4% paraformaldehyde/PBS using Rhodamine RedX (RRX)-conjugated secondary antibodies anti-rabbit and Fluorescein (FITC)-conjugated secondary antibodies anti-mouse (Jackson ImmunoResearch).

For *in vitro* differentiated adipocytes, the stromal vascular fraction (SVF) was isolated from brown adipose tissue depots after collagenase type I/dispase II digestion in 4% BSA KRH buffer for 45 minutes then washed, filtered and spun down at 900 rpm for 10 minutes. Cells were cultured in high glucose DMEM with 10% fetal bovine serum (Hyclone) and 1X pen/strep until confluence. Two days later, *in vitro* adipogenic differentiation was induced using standard insulin, IBMX (Sigma), DEX (Sigma) and T3 (Sigma) cocktail for 2 days then changed to maintenance media containing high glucose DMEM with 10% fetal bovine serum, pen/strep, and insulin for 12 additional days before being fixed in 10% formalin to perform Oil Red O staining as described^6^.

**Protein extraction, Separate extracts preparation, Submitochondrial Localization, *In Vivo* Sumoylation assay and Immunoprecipitation**. For whole cell extracts preparation, cells were rinsed in PBS, harvested and incubated for 20’ on ice in IPH buffer (50 mM Tris-HCl pH 8.0, 150 mM NaCl, 5 mM EDTA, 0.5% NP-40, 50 mM NaF,2 mM Na_2_VO_3_, 1mM PMSF and protease inhibitor mix). For cytoplasmatic, mitochondrial and nuclear extracts fractionation cells were rinsed in PBS, harvested and re-suspended in gradient buffer (10 mM HEPES pH 7.9, 1mM EDTA, 210 mM Mannitol, 70mM Sucrose, 10mM NEM, 50 mM NaF, 2 mM Na_2_VO_3_, 1mM PMSF and protease inhibitors cocktail) then homogenized via 10 passages through 25G syringe followed by low-speed centrifugation for 10 min. The nuclear pellet was incubated for 20 min in high-salt buffer (10 mM Hepes pH 7.9, 20% glycerol, 420 mM NaCl, 1.5 mM MgCl_2_, 0.2mM EDTA, 0.5mM DTT, 10mM NEM, 50 mM NaF, 2 mM Na_2_VO_3_, 1mM PMSF and protease inhibitor mix) while the supernatant was recovered and subjected to high-speed centrifugation to separate the mitochondrial pellet from the cytoplasmic fraction. The mitochondrial pellet was incubated for 15 min in lysis buffer (50 mM Tris/HCl pH 8, 300 mM NaCl, 1mM EDTA, 1% Triton X-100, 10mM NEM, 50 mM NaF, 2 mM Na_2_VO_3_,1mM PMSF and protease inhibitor mix). To examine submitochondrial localization, the isolated mitochondria fraction was treat with either Proteinase K (2, 5 or 10 ng) in ice for 30 min or 50 **μ**g/ml trypsin in ice for 30 min under either isotonic or hypotonic condition, reaction was terminated adding respectively 1mM PMSF or 10% TCA. Concentration of protein extracts was measured using the colorimetric BIORAD assay. Extracts were boiled in SDS sample buffer and loaded 10% Mini-PROTEAN TGX gels (Biorad), prior to transfer onto PVDF membranes (Millipore) and western blotting following standard protocols. For *in vivo* Sumoylation assay cells were transfected with HA-SUMO1 and or either HA-GPS2-Flag full length or HA-GPS2-Flag K45,75R then cytosolic, mitochondrial and nuclear extracts were separated by electrophoresis or subjected to immunoprecipitation as previously described^7^.

### RNA Isolation, RT-PCR Analysis and RNA-seq

RNA was isolated form cell or mice tissue following the manufacturer protocol for the RNeasy Kit (QIAGEN). First strand cDNA synthesis from total RNA template was performed using the Biorad IScript cDNA Synthesis System, followed by SYBR-green qPCR amplification. Normalization was performed using specific amplification of *CyclophilinA* and qPCRs were performed in triplicate for each biological experiment. Data are shown as sample mean between triplicate experiments +/-standard deviation. Significance was calculated by paired student’s T-test. Primers sequences used for each specific genes are available on request. For the RNA-seq, cells were subjected to standard RNA isolation prior to RNA library preparation following Illumina’s RNA-Seq Sample Preparation Protocol. Resulting cDNA libraries were sequenced on the Illumina’s HiSeq 2000.

### ChIP assay and ChIP-seq

Chromatin immunoprecipitation (ChIP) was performed as described^2^. Briefly, approximately 10^7^ cells were cross-linked with 1% formaldehyde at room temperature (~25 °C) for 10 min and neutralized with 0.125 M glycine. When Brown Tissue was used, tissue was minced with curved scissors (slurry like) and put in 10ml tubes containing PBS 1% formaldehyde for crosslink for 15 min at RT and neutralized with 0.125 M glycine. Tissue was then shredded in Lysis Buffer using a Bullet Blender Homogenizer at max power for 5 min 4°C. After sonication, chromatin was incubated with 2μg of antibody at 4 °C overnight. Immunoprecipitated complexes were collected using sepharose beads A (Life Technologies). After extensive washes the DNA was extracted and purified by phenol/chloroform extraction. ChIP experiments were repeated at least three times and representative results are shown as samples mean between technical replicates +/-standard deviation. Significance was calculated by paired student’s T-test. Primers used are specific for the regions tested and their sequences are available on request. For ChIP-Seq, the extracted DNA was ligated to specific adaptors followed by deep sequencing on the Illumina’s HiSeq 2000 according to the manufacturer’s instructions.

### Gro-seq assay

GRO-seq experiments were performed as previously reported^8^. Briefly, 3T3-L1 cells were swelled in swelling buffer (10mM Tris-Cl pH 7.5, 2mM MgCl_2_, 3mM CaCl_2_) for 5min on ice and then lysed in lysis buffer (swelling buffer with 0.5% IGEPAL and 10% glycerol, 4U/ml of superase in) in ice for 5 min then nuclei were pellet and re-suspended in 100μl of freezing buffer (50mM Tris-Cl pH 8.3, 40% glycerol, 5mM MgCl_2_, 0.1mM EDTA). For the run-on assay, re-suspended nuclei were mixed with an equal volume of reaction buffer (10mM Tris-Cl pH 8.0, 5mM MgCl_2_, 1mM dithiothreitol (DTT), 300mM KCl, 20 units of Superase-In, 1% sarkosyl, 500μM ATP, GTP, Br-UTP and 2μM CTP) and incubated for 5min at 30°C. The nuclear-run-on RNA (NRO-RNA) was then extracted with TRIzol LS reagent (Invitogen) following the manufacturer’s instructions. After base hydrolysis on ice for 40min and followed by treatment with DNase I and antarctic phosphatase, the Br-UTP-labelled NRO-RNA was purified by anti-BrdU argarose beads (Santa Cruz Biotech) in binding buffer (0.5× SSPE, 1mM EDTA, 0.05% Tween) for 3h at 4°C while rotating. Then T4 PNK (NEB) was used to repair the end of NRO-RNA. Subsequently, complementary DNA synthesis was performed as reported^8^ with few modifications. The RNA fragments were subjected to the poly-A-tailing reaction by poly-A polymerase (NEB) for 30min at 37°C. Reverse transcription was then performed using superscript III (Invitrogen) with oNTI223 primer (available on request). The cDNA products were separated on a 10% polyacrylamide TBE-urea gel with the right product (~100–500bp) being excised and recovered by gel extraction. After that, the first-strand cDNA was circularized by CircLigase (Epicentre) and re-linearized by Ape1 (NEB). Re-linearized single-strand cDNA was separated by TBE gel and the products of the desired size were excised (~120–320bp) for gel extraction. Finally, the cDNA template was amplified by PCR using the Phusion High-Fidelity enzyme (NEB) with primers oNTI200 and oNTI201 for deep sequencing (available on request) on the Illumina’s HiSeq 2000 system according to the manufacturer’s instructions.

### Bioinformatics analysis of ChIP-seq, RNA-seq and GRO-seq datasets

For the bioinformatics analyses, we have build comprehensive lists of mouse and human mitochondrial genes by combining the sets of mitochondrial genes from MITOMINER database^9^ with the sets of genes that are annotated as mitochondrial from NCBI, AMIGO, and ENSEMBL databases. The ChIP-seq bioinformatics analyses were performed as previously described^3,7^. The GPS2 ChIP-seq datasets were downloaded from NCBI GEO database (GSE35197^3^, GSE57777^7^ and GSE66774^10^), PPAγγ, RXR, and POL2 ChIP-seq datasets were downloaded from NCBI GEO series GSE13511^11^, NCoR and SMRT ChIP-seq datasets were downloaded from ArrayExpress E-MTAB-103^12^. The alignment of POL2 and H3K9me3 ChIP-seq samples was performed by using Bowtie2^13^ to mm8 assembly of the mouse genome, and sets of equal number of reads were randomly extracted from siCTL and siGPS2 aligned samples. We have normalized the ChIP-seq datasets on specific genomics regions by using TMM normalization procedure in edgeR^14^, and the normalized counts were obtained by using cpm() function. HOMER software suite^15^ was used to call the ChIP-seq peaks and to compute the heatmaps, the read density profiles, the motif density profiles on promoters, and to find the enriched motifs. The heatmaps of ChIP-seq datasets were displayed by using R packages pheatmap and gplots, and the boxplots were displayed in ggplot2.

The RNA-seq fastq files (siCTL, siGPS2) were aligned by using Tophat^16^ and the sequencing reads were counted over transcripts by using Cufflinks^17^, FeatureCounts^18^ and HOMER^15^. Differential expressed genes across two independent RNAseq replicate experiments were identified by edgeR^14^ using FDR<0.05. GO/pathway enrichment was computed with DAVID^19,20^, ToppFun, and in enrichR^21,22^. The boxplots and scatterplots were displayed in R standard libraries, in ggplot2, and in limma.

The GRO-seq fastq files (siCTL, siGPS2) were aligned to mouse genome mm8 by using bowtie2; we have used the same number (4.2 millions) of randomly extracted reads for computing the gene expression levels. The aligned reads were counted over the RefSeq gene bodies (after excluding a TSS 400bp-proximal region, and TTS 400bp-distal region) by using bedtools^23^. The statistically significant differentially expressed genes were defined by edgeR^14^ for a BCV of 0.01. We have also used a GRO-seq dataset from NCBI GEO (GSE56747^24^) for computing the read density profiles on TSSes. All our dataset are available in NCBI GEO (GSE80994).

### Immunogold-Electron Microscopy

For preparation of cryosections cells were rinsed in PBS, harvested and fixed in 4% paraformaldehyde/PBS for two hour then cells were infiltrated with 2.3M sucrose/PBS/glycin 0.2M) and freezed in liquid nitrogen. Frozen samples were sectioned at −120°C and transferred to formvar-carbon coated copper grids and goaldlabeling wad carried out following standard protocols (Harvard EM Core).

### Mitochondrial content

Total DNA was extracted from cells using QuickExtract DNA Extraction Solution 1.0 (Epicenter) following manufacturer’s instructions. DNA amplification of the mitochondrial-encoded NADH dehydrogenase 1 (mt-ND1) relative to nuclear TFAM was used to determine mitochondrial DNA copy numbers.

### Mitochondrial Isolation and Mass

Brown Adipose Tissue was isolated from GPS2-WT and KO mouse and weighed. Isolated BAT was rinsed and minced in ice-cold PBS then homogenized in a glass-teflon dounce homogenizer containing SHE pH=7.2 (250 mM Sucrose, 5mM HEPES, 2 mM EGTA, BSA 2%.) + BSA buffer. After 9-10 strokes throught the tefflon pestle the homogenate was centrifuged at 900xg for 10 min at 4°C. The resulting supernatant was then centrifuged at 9000xg for 10 min at 4°C and the pellet was washed once and then re-suspended in SHE without BSA. Protein content was measured by BCA. Mitochondiral Mass was calculated as ratio between protein concentrations expressed in μg and tissue weight expressed in mg.

### Respirometry

Cells were plated in Seahorse V.7 multi-well culture plates. The day after, media was replaced by running media (XF Seahorse Assay Media supplemented with 5.5mM glucose, 0.5mM pyruvate and 1mM glutamine) and the plate was placed at 37°C for 1 h (no carbon dioxide). Oxygen consumption was measured at 37°C using a Seahorse XF24 Extracellular Flux Analyzer (Seahorse Bioscience, Billerica, MA). Mitochondrial stress test compounds (10 μM oligomycin, 2.5 μM FCCP and 10 μM antimycin A) were injected through ports A, B, and C, respectively, to measure mitochondrial respiration linked to ATP synthesis, leak, maximal respiratory capacity and non-mitochondrial oxygen consumption according to the manufacturer’s instructions.

For mitochondrial activity, 4μg of mitochondrial protein fractions were loaded per well for complex I driven respiration (pyruvate+malate) and 2μg for complex II-driven respiration (succinate+rotenone) in 25 μl of Mitochondrial Assay Solution pH 7.2 (MAS: 100mM KCl, 10mM KH2PO4, 2mM MgCl2, 5mM HEPES, 1mM EGTA, 0.1% BSA and 1mM GDP) per Seahorse XF96 well. The plate was centrifuged at 4°C, 5 min at 3400 rpm. Then, 110 μl of MAS with the respective fuels were carefully added per well. Plate was warmed at 37°C for 4 min then respirometry assay within the XF96 was performed as described above for the XF24. Pyruvate was used at 5mM, malate 5mM, succinate 5mM and rotenone 2μM. ADP was injected at port A (3.5mM), Oligomycin A at port B (3.5μM), FCCP at port C (4μM) and Antimycin A at port D (4μM).

## Supplemental Information

### GPS2 regulates mitochondria biogenesis via mitochondrial retrograde signaling and chromatin remodeling of nuclear-encoded mitochondrial genes

Maria Dafne Cardamone, Bogdan Tanasa, Carly Cederquist, Jiawen Huang, Kiana Mahdaviani, Wenbo Li, Michael G. Rosenfeld, Marc Liesa and Valentina Perissi

**Figure S1.**
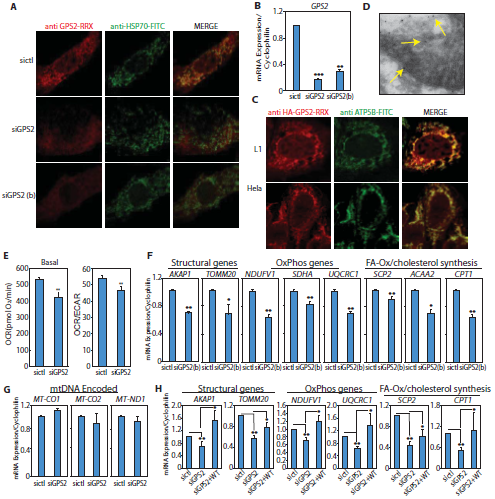
Related to Figure 1. **(A)** Mitochondria localization of endogenous GPS2 by IHF staining in murine 3T3-L1 cells, with two different siRNA against GPS2 used to confirm signal specificity. Co-staining of endogenous mtHSP70 serves as a mitochondrial marker. **(B)** siRNA efficiency measured by qPCR analysis in transiently transfected 3T3-L1 cells. **(C)** Mitochondria localization of overexpressed HA-GPS2 by IHF staining in murine 3T3-L1 and human Hela cells. Co-staining of endogenous mtATP5B serves as a mitochondrial marker. **(D)** Mitochondria labeling by Immunogold EM for GPS2 in Hela cells. **(E)** Decreased basal respiration and bioenergetic profile in Hela cells upon GPS2 downregulation. (**F)** RT-qPCR analysis of different classes of neMITO genes expression in 3T3-L1 cells transfected with siCTL or siGPS2(b). **(G)** RT-qPCR analysis of the expression of representative mitochondria-encoded genes in 3T3-L1 cells transfected with siCTL or siGPS2. **(H)** Rescued expression of representative neMITO genes in GPS2-depleted 3T3-L1 cells by overexpressing HA-GPS2. Results in **(B), (E), (F), (G)** and **(H)** represent the sample mean of three independent experiments; error bars represent s.e.m. (n=3). *indicate p-value<0.05; *indicate p-value<0.01.

**Figure S2.**
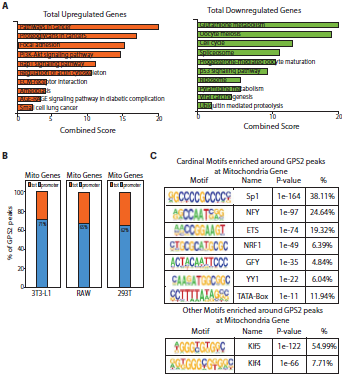
Related to Figure 2. **(A)** DEGs between WT and GPS2-KD 3T3-L1 cells by RNAseq. Analysis of the 10 most significant pathways associated with total upregulated (left) and downregulated (right) genes by EnrichR (KEGGPathways). **(B)** Percentage of GPS2 peaks on mitochondrial genes localized to promoter areas (+1kb/-200bp around TSS) in ChIPseq data sets from 3T3-L1 (GSE57779), 293T (GSE35197) and RAW (GSE66774) cells. **(C)** Cardinal DNA-binding motifs and other TFs binding motifs enriched around the GPS2 peaks in 3T3-L1 cells as calculated by HOMER Known Motifs enrichment analysis.

**Figure S3.**
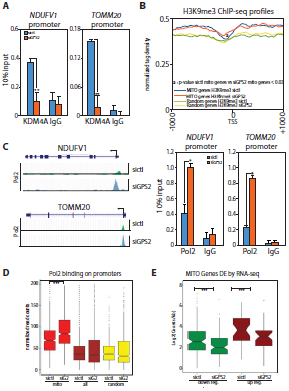
Related to Figure 3. **(A).**Decrease in JMJD2A/KDM4A recruitment to *Ndufv1* and *Tomm20* gene promoters by ChIP in GPS2-KD 3T3-L1 cells. **(B)** Tag density plot showing the increase in H3K9me3 binding on mitochondrial genes in GPS2-KD 3T3-L1 cells compared to randomly selected groups of comparable size (2086 genes). P-values were computed using the Wilcoxon test based on the difference in normalized counts +/-200bp around the TSS. **(C)** Representative ChIPseq tracks for POL2 binding to *Ndufv1* and *Tomm20* gene promoters on the UCSC browser. Validation of POL2 accumulation on target promoters in GPS2-KD 3T3-L1 cells by hand-ChIP. **(D)** Boxplots showing a significant increase in POL2 binding specific to the promoters of mitochondrial genes in GPS2-KD 3T3-L1 cells. P-values were computed using the Wilcoxon test based on the difference in normalized counts +/-200bp around the TSS (Replicate 1, p-value < 2.2e-16; Replicate 2 p-value<2.99e-09). **(E)** Boxplot showing decreased transcription by GROseq for both down-and up-regulated mitochondrial genes as defined by RNAseq in 3T3-L1 transfected with siCTL or siGPS2 (p-value<2.2e-16 for both groups). Hand-ChIP data are representative of three independent experiments, bar graphs represent the sample mean of three technical replicates +/-SD. *indicate p-value<0.05; *indicate p-value<0.01, *indicate p-value<0.001.

**Figure S4.**
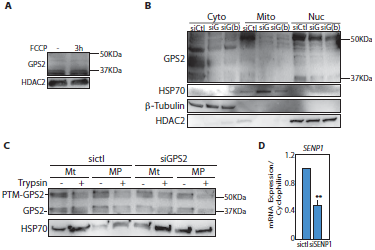
Related to Figure 4. **(A)** Total GPS2 protein levels not changing upon FCCP treatment as shown by WB of whole cell extracts. **(B)** Two different siRNA against GPS2 -siG and siG(b)-were used to show by western blotting of fractionated extracts the specificity of GPS2 bands. **(C)** Trypsin assay showing localization of GPS2 and post translational modified GPS2 (PTM-GPS2) in the mitochondria matrix and outer membrane respectively. Mt stands for isolated mitochondria, MP for mitoplasts. **(D)** SENP1 siRNA efficiency by qPCR analysis. Results represent the sample mean of three independent experiments; error bars represent s.e.m. (n=3). ^*^ indicate p-value<0.05; ^**^ indicate p-value<0.01.

**Figure S5.**
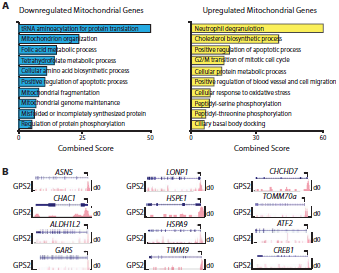
Related to Figure 5. Analysis of 10 most enriched pathways among mitochondrial genes upregulated or downregulated in response to FCCP depolarization as identified by GROseq (EnrichR, KEGG Pathways). **(B)** Representative ChIPseq tracks on the UCSC browser for GPS2 binding in 3T3-L1 cells to genes representative of the pathways indicated in Figure 5C.

